# Mitochondrial fusion controls the development of specialized mitochondrial structure and metabolism in rod photoreceptor cells

**DOI:** 10.1101/2025.05.21.655403

**Authors:** Michael Landowski, Ryo Hagimori, Purnima Gogoi, Vijesh J. Bhute, Sakae Ikeda, Tetsuya Takimoto, Akihiro Ikeda

## Abstract

Mitochondria are dynamic organelles that undergo continuous morphological changes, yet exhibit unique, cell-type-specific structures. In rod photoreceptor cells of the retina, these structures include elongated mitochondria in the inner segments and a distinct, large, circular mitochondrion in each presynaptic terminal. The mechanisms underlying the establishment and maintenance of these specialized mitochondrial morphologies, along with their functional significance, are not well understood. Here, we investigate the roles of mitochondrial fusion proteins mitofusin 1 (MFN1) and mitofusin 2 (MFN2) in shaping these structures and maintaining photoreceptor cell health. Rod photoreceptor cell-specific ablation of MFN1 and MFN2 resulted in mitochondrial fragmentation by one month of age, suggesting that mitochondrial fusion is essential for the development of photoreceptor cell-specific mitochondrial structures. Notably, the layer structures of the retina examined by light microscopy appeared unaffected at this age. Following this time period, significant photoreceptor cell degeneration occurred by three months of age. Furthermore, we showed that impaired mitochondrial fusion perturbed the balance of proteins involved in glycolysis, oxidative phosphorylation (OXPHOS), and β-oxidation, highlighting the critical role of mitochondrial fusion in ensuring the proper levels of proteins necessary for optimal energy metabolism. Additionally, we identified upregulation of cellular stress pathways such as endoplasmic reticulum (ER) stress and unfolded protein response (UPR), which arise in response to energy deprivation, and cytoprotective biosynthetic pathways mediated by CCAAT/enhancer-binding protein gamma (C/EBPγ) and mammalian target of rapamycin complex 1 (mTORC1) signaling. In summary, our findings indicate that mitochondrial fusion through MFN1 and MFN2 is vital for the development of unique mitochondrial structures and proper energy production, underscoring the fundamental importance of mitochondrial dynamics in photoreceptor cell function and survival.

**Significance Statements:** Rod photoreceptor cells exhibit unique mitochondrial morphologies and high energy requirements. In this report, we examined how these unique mitochondrial structures are established and their biological significance. We identified that mitochondrial fusion is essential for the development of characteristic mitochondrial morphologies in rod photoreceptor cells. Furthermore, we demonstrated that impaired mitochondrial fusion disrupts the equilibrium of proteins associated with OXPHOS, glycolysis, and β-oxidation, ultimately leading to an imbalance in cellular energy homeostasis. Our findings also revealed activation of cellular stress pathways, including ER stress and the UPR, which are likely triggered by energy depletion. Additionally, we identified activation of cytoprotective biosynthetic pathways that are engaged to preserve cellular homeostasis and function.

## Introduction

Mitochondria, the energy-generating organelles, dynamically fuse and fission to take various forms in cells. Mitochondrial structure is generally dynamic in nature, and some cells display very unique mitochondrial structures. However, the exact relations between their unique form and associated function are still not fully understood. Retinal rod photoreceptor cells, the energy-intensive neurons, provide an excellent model for investigating the significance of mitochondrial form and its function since they exhibit a uniquely uniform arrangement of elongated mitochondria in the inner segments and one large circular mitochondrion in each of the presynaptic terminals (1–3).

Energy homeostasis in rod photoreceptor cells is unique in that they use more than 80% of glucose for aerobic glycolysis converting it to lactate, rather than complete respiration including glycolysis, tricarboxylic acid (TCA) cycle, and oxidative phosphorylation (OXPHOS) (4, 5). Nonetheless, they do rely on OXPHOS for energy production since less than 20% of glucose that enters OXPHOS has been proposed to account for 80% of the total ATP generation (5). Energy production in rod photoreceptor cells occurs mainly in the inner segments (6), where the glycolytic system is dominant (5, 7) and numerous characteristically elongated mitochondria are present (1, 3)Synapses also require large amounts of energy to circulate neurotransmitters, which is maintained by local OXPHOS and glycolysis (8). While mitochondria in rod photoreceptor cells show unique localization and morphology, it is still not completely clear how mitochondria regulate energy-generating systems such as glycolysis and OXPHOS, and contribute to the homeostasis of rod photoreceptor cells.

Mitochondrial dynamics have been linked to complex cellular processes such as metabolism, immune response, and cell death. Mitochondria maintain ATP-producing capacity and homeostasis through fission and fusion, and the balance of mitochondrial fission and fusion is tightly regulated in accordance with cellular metabolic states (9). Impaired mitochondrial dynamics cause energy disruption, and ultimately lead to cell death (10, 11). Mitochondrial fusion is stimulated by high-energy demand to maximize the energy production (9). Mitofusin (MFN) 1 and MFN2 contribute to mitochondria fusion and regulate proper mitochondrial dynamics (12, 13). Studies have shown that MFN1 or MFN2 deficiency results in abnormal energy production and defective biosynthetic processes (14). For example, it has become clear using gene targeted mice that mitochondrial fusion in pro-opiomelanocortin neurons regulates intracellular metabolism and maintains its function robustly (15, 16). However, the role of mitochondrial dynamics in photoreceptor cell-specific structures and metabolic activities is still to be determined.

In this study, we ablated MFN1 and MFN2 specifically in rod photoreceptor cells and observed mitochondrial fragmentation, suggesting that proper mitochondrial fusion is necessary for establishing rod photoreceptor cell-specific mitochondrial structures. Subsequent to mitochondrial fragmentation, we observed significant photoreceptor cell degeneration. We demonstrated that impaired mitochondrial fusion disturbed the levels of proteins involved in OXPHOS and mitochondrial β-oxidation, suggesting a pivotal role for mitochondrial fusion in regulating efficient energy metabolism in rod photoreceptor cells. Additionally, our multi-omics analysis revealed that energy disruption in these cells can activate cellular stress pathways and cytoprotective mechanisms. In summary, our study suggests that proper mitochondrial morphologies in rod photoreceptor cells maintained through mitochondrial fusion is crucial for sustained and optimized energy production as well as maintenance of the cell integrity.

## Results

### MFN1 and MFN2 Are Necessary for Development of Rod-Specific Mitochondrial Morphology and Integrity of Rod Photoreceptor Cells

We ablated genes encoding mitochondrial fusion factors, MFN1and MFN2, in murine rod photoreceptor cells by crossing mice with floxed alleles of *Mfn1* (*Mfn1^flx^*, (17)) and *Mfn2* (*Mfn2^flx^*,(17)) and *Rho-iCre* (*Rho-Cre*) transgenic mice, in which Cre-recombinase is expressed specifically in rod photoreceptor cells (17)(Supplementary Fig. 1A). We confirmed reduction of MFN1 and MFN2 levels by approximately 10–15% in the neural retina of these mice, where MFN1 and MFN2 were still expressed in other retinal cells (Supplementary Fig. 1B). To investigate the role of mitochondrial fusion in shaping the unique mitochondrial structures and function in rod photoreceptor cells, we examined mitochondrial morphologies and retinal health in mice with rod-specific deletion of MFN1 and MFN2 (*Rho-Cre/Mfn1^flx/flx^/Mfn2^flx/flx^*). Electron microscopy (EM) revealed significantly increased mitochondrial fragmentation in the inner segments of rod photoreceptor cells in *Rho-Cre/Mfn1^flx/flx^/Mfn2^flx/flx^* mice (Fig. 1A, B). To further characterize mitochondrial morphology within the inner segment, we conducted three-dimensional confocal imaging using a mitochondrial marker, TOMM20. Wild-type (WT) mice appeared to contain predominantly elongated mitochondria in the inner segment, whereas *Rho-Cre/Mfn1^flx/flx^/Mfn2^flx/flx^* mice displayed a marked increase in fragmented mitochondrial structures within the same region. (Fig. 1C). EM analysis revealed significantly increased mitochondrial fragmentation in the synapse of rod photoreceptor cells in *Rho-Cre/Mfn1^flx/flx^/Mfn2^flx/flx^* mice (Fig. 1D, E), accompanied by a significant decrease in mitochondrial size within the synapse (Fig 1F). While *Rho-Cre/Mfn1^flx/flx^/Mfn2^+/+^* mice and *Rho-Cre/Mfn1^flx/+^/Mfn2^flx/flx^* mice exhibit intact mitochondrial architecture, *Rho-Cre/Mfn1^flx/flx^/Mfn2^flx/+^* mice showed moderately fragmented mitochondria in the synapse and inner segments (Fig. 1A, B, D, E). No changes in mitochondrial aspect ratio were observed in synapses of any of the genotypes (Supplemental Fig. 1C). These findings highlight the crucial role of MFN1 and MFN2-mediated mitochondrial fusion in shaping the distinct mitochondrial architectures within rod photoreceptor cells.

**Figure 1.**
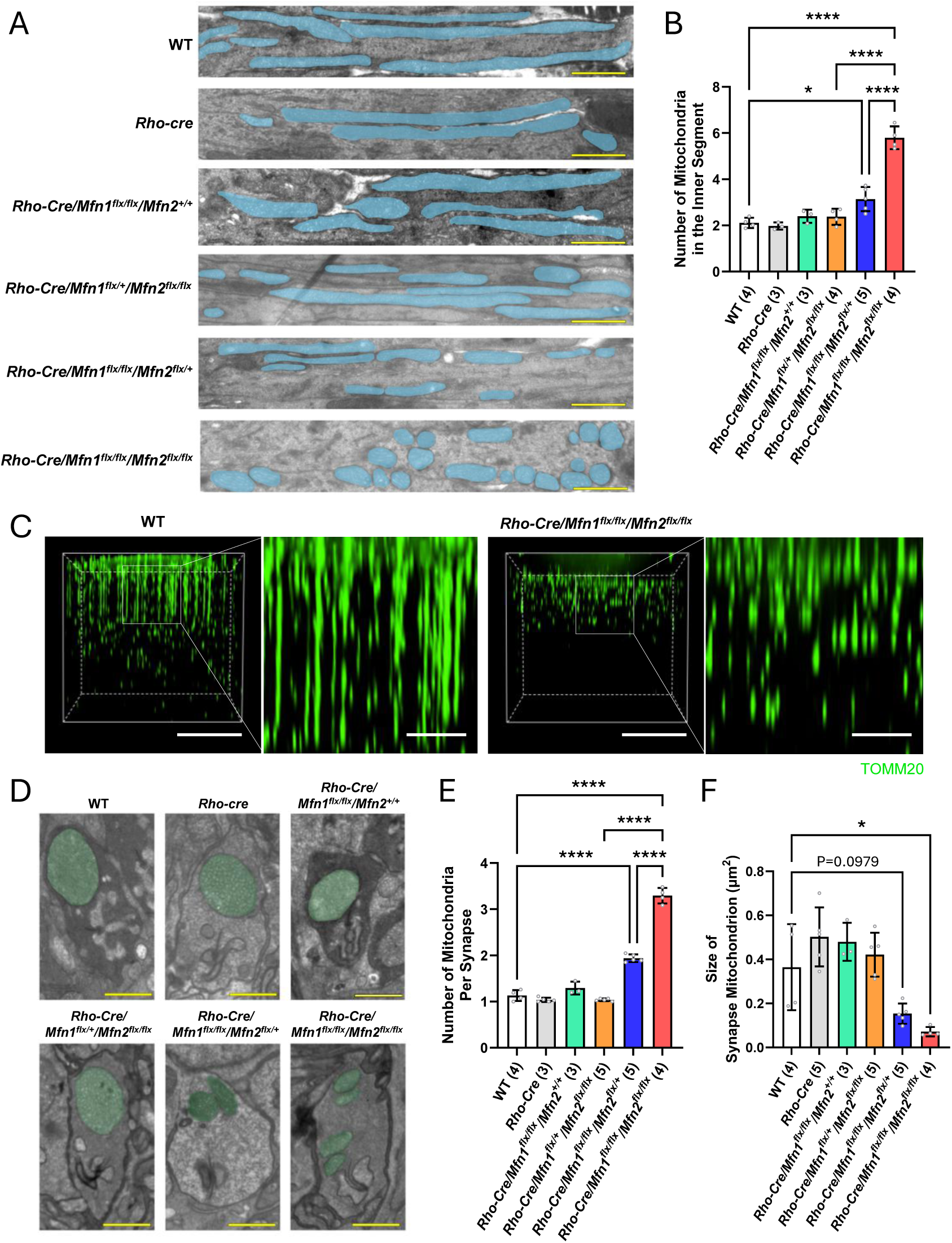
Abnormal mitochondrial morphologies in rod photoreceptor inner segments and synapses due to ablation of *Mfn1* and *Mfn2*. (A) Representative electron micrographs of rod photoreceptor inner segments. Mitochondria are shaded in blue. Magnification = 8,800X. Scale bar = 1 micron. (B) Quantification of the number of mitochondria in rod photoreceptor inner segments. (C) Representative three-dimensional reconstructions of mitochondria within photoreceptor inner segments, visualized by immunostaining with TOMM20. Scale bar = 20 μm; scale bar for magnified view = 5 μm. (D) Representative electron micrographs of rod photoreceptor synapses. Mitochondria are shaded in green. Magnification = 8,800X. Scale bar = 1 micron. (E) Quantification of the number of mitochondria in the rod photoreceptor synapse. (F) Quantification of the size of mitochondria in the rod photoreceptor synapse. Dots represent individual data points. Number in the parenthesis denotes the number of mice used in the experiment. Data is presented as mean +/-SD. *P<0.05, ****P<0.0001 by two-way ANOVA with post-hoc Tukey’s test.

Histological analysis at the light microscopy level showed no gross abnormalities in rod photoreceptor cells at one month of age in any of the genotypes (Fig. 2A and B). However, by three months of age, significant photoreceptor cell degeneration was observed in *Rho-Cre/Mfn1^flx/flx^/Mfn2^flx/flx^* mice (Fig. 2C and D). This indicates that proper mitochondrial structures, facilitated by mitochondrial fusion, are critical for maintaining the integrity of rod photoreceptor cells.

**Figure 2.**
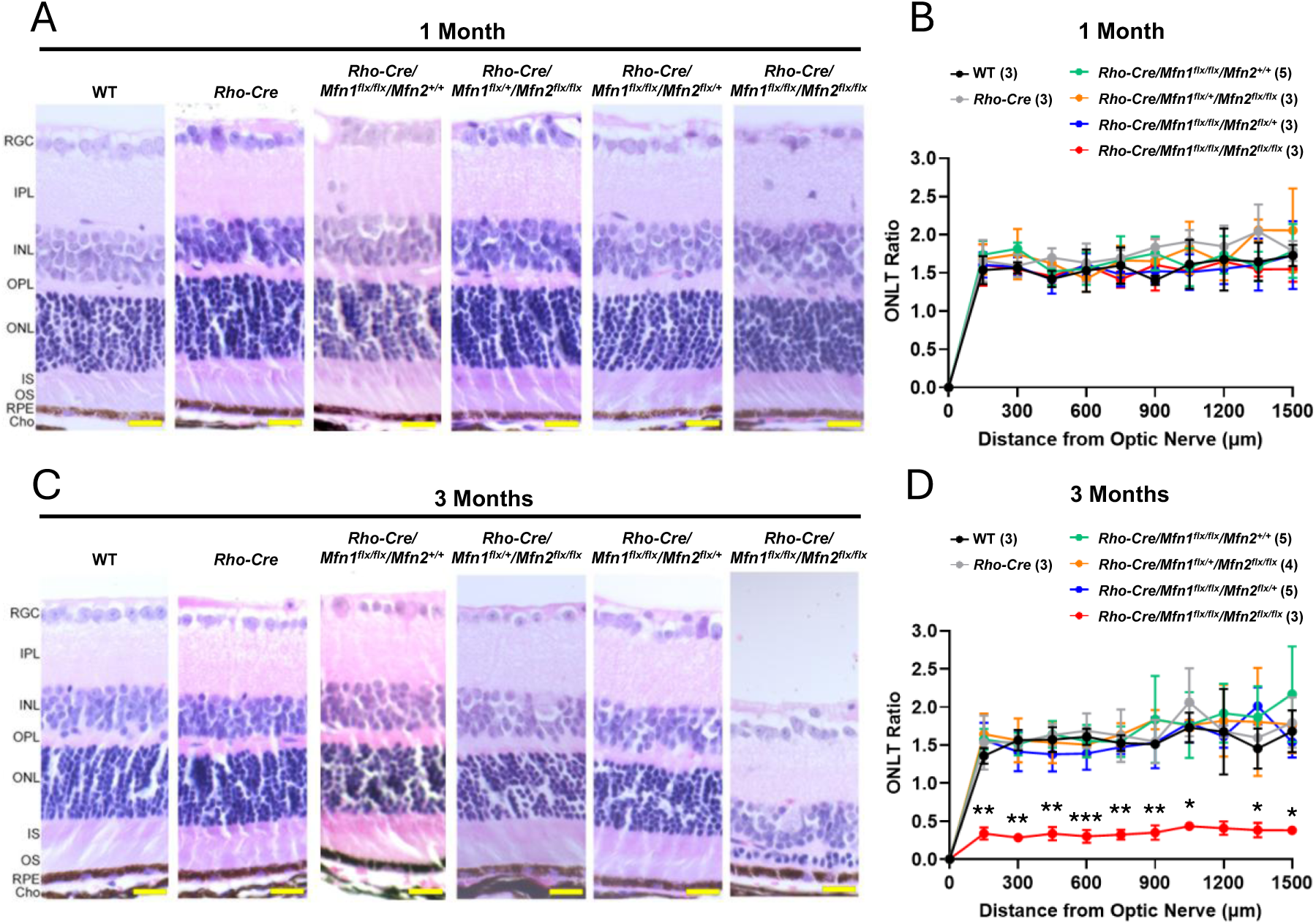
Photoreceptor cell degeneration in mice with rod-specific ablation of *Mfn1* and *Mfn2*. (A) Representative images of H&E-stained retinal sections of one-month-old mice. Magnification = 40X. Scale bar = 20 microns. (B) Outer nuclear layer thickness (ONLT) ratios. (C) Representative images of H&E-stained retinal sections of three-month-old mice. Magnification = 40X. Scale bar = 20 microns. (D) ONLT ratios. Note that *Rho-Cre/Mfn1^flx/flx^/Mfn2^flx/flx^* mice exhibit ONL degeneration at three months of age. Number in the parenthesis denotes the number of mice used in the study. Data is presented as mean +/-SD. *P<0.05, **P<0.01, ***P<0.001, ****P<0.0001 by two-way ANOVA with post-hoc Tukey’s test.

### Molecular Pathways Altered by Ablation of *Mfn1* and *Mfn2*

To elucidate the molecular pathways associated with rod photoreceptor cell degeneration in *Rho-cre/Mfn1^flx/flx^/Mfn2 ^flx/flx^* mice, we performed RNA sequencing (RNA-seq) on neural retinas collected from one-month-old WT and *Rho-cre/Mfn1^flx/flx^/Mfn2 ^flx/flx^* mice before degeneration started. Differential expression analysis identified 974 dysregulated genes (399 upregulated and 575 downregulated) in *Rho-cre/Mfn1^flx/flx^/Mfn2 ^flx/flx^* mice compared to WT controls, indicating significant transcriptional alterations.

In the neural retina of *Rho-cre/Mfn1^flx/flx^/Mfn2 ^flx/flx^* mice, cell-specific markers (total 75) are not significantly changed or showed log2 fold changes (LogFC) <1 overall except for 9 markers of macrophage, Müller cell, and oligodendrocyte (Supplementary Fig. 2), suggesting that changes in the cellular composition were minimal. This finding was consistent with the results of histological analysis by light microscopy (Fig. 2A).

To investigate the functional relevance of dysregulated genes, we conducted gene set enrichment analysis (GSEA) to identify the top 10 enriched gene sets, using LogFC values between WT and *Rho-cre/Mfn1^flx/flx^/Mfn2 ^flx/flx^* (conditional KO [cKO]) retinas with the fast gene set enrichment analysis algorithm (Fig.3A). A positive normalized enrichment score (NES) indicated that genes upregulated in cKO retinas were predominantly enriched in the top-ranking gene sets. Pathway analysis revealed that genes involved in ‘unfold protein response (UPR)’ and ‘CCAAT/enhancer-binding protein gamma (C/EBPγ)’ (Fig. 3A), both of which are known to respond to ER stress (18–20) were enriched. Furthermore, overrepresentation analysis of the genes in CEBPG_TARGET_GENES with the highest expression variation ratios confirmed the enrichment of genes related to amino acids (AA) metabolism and translation process in *Rho-cre/Mfn1^flx/flx^/Mfn2 ^flx/flx^* mice (Fig. 3B). Our GSEA results also revealed that the gene set involved in mammalian target of rapamycin complex 1 (mTORC1) signaling was enriched to the higher level of the gene set expression variability ratio in *Rho-cre/Mfn1^flx/flx^/Mfn2 ^flx/flx^* mice (Fig. 3C). Notably, in addition to these findings, several key glycolytic genes were suppressed, indicating a shift away from glycolytic energy production in rod photoreceptor cells of *Rho-cre/Mfn1^flx/flx^/Mfn2 ^flx/flx^* mice (Fig. 3C).

**Figure 3.**
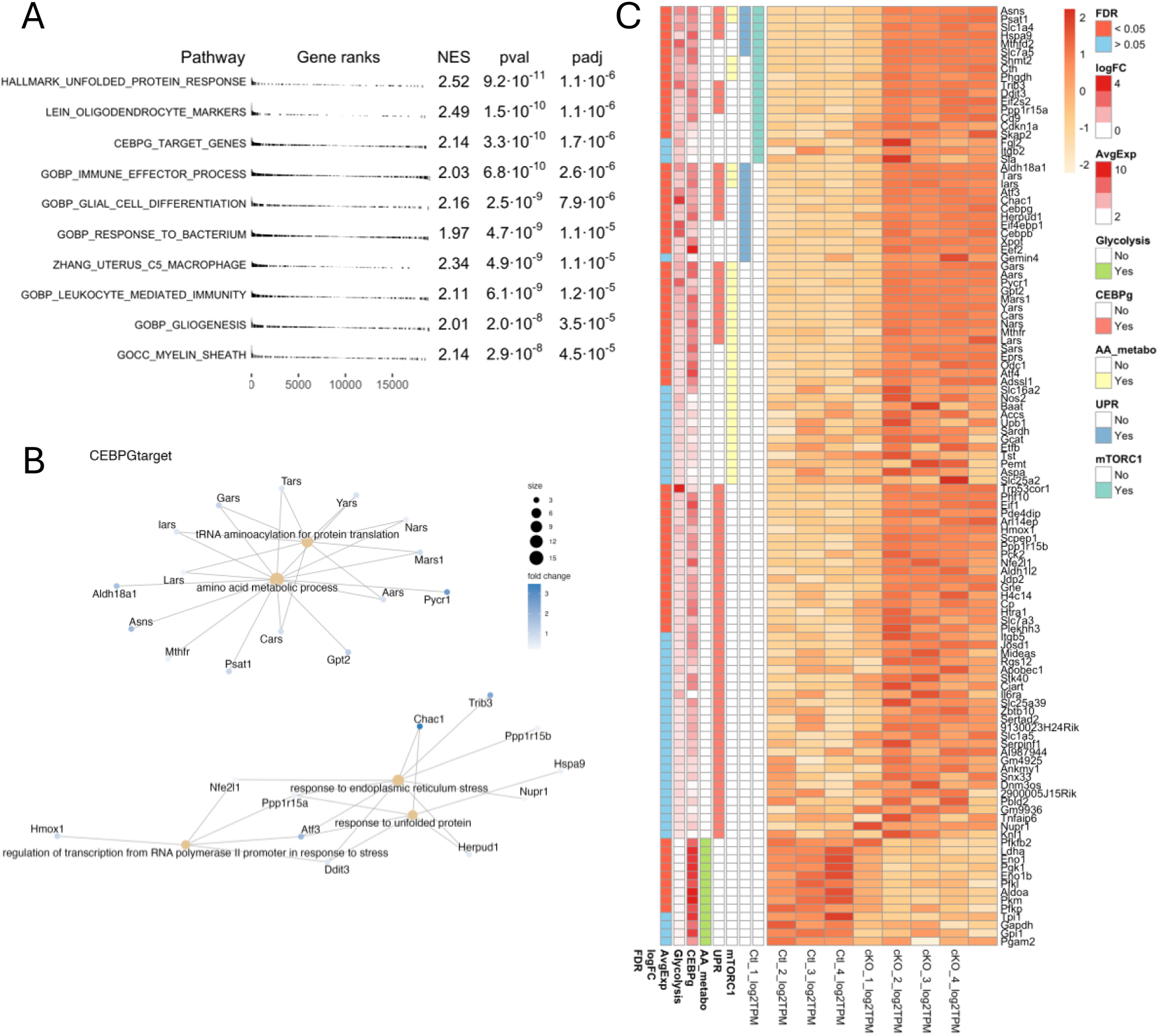
**Molecular pathways altered in the neural retina by rod-specific ablation of *Mfn1* and *Mfn2*** (A) Top10 enriched pathways were identified by Gene Set Enrichment Analysis (GSEA) and ranked by P value (pval) and adjusted P value (padj). NES stands for normalized enrichment score. (B) Overrepresentation analysis of the CEBPG_TARGET_GENES gene sets with the highest expression variation ratio. (C) Heatmap showing genes involved in mTORC1 signaling, amino acid (AA) metabolism, unfold protein response (UPR), and C/EBPγ that are significantly enriched, and glycolysis-related genes that are significantly downregulated in the neural retina by rod-specific ablation of *Mfn1* and *Mfn2*. False discovery rate (FDR), logFC, and average expression (AvgExpr) are shown in the left column for each gene.

### Metabolomic Changes Resulting from Ablation of *Mfn1* and *Mfn2*

To investigate the difference of metabolites influenced by mitochondrial fragmentation in *Rho-cre/Mfn1^flx/flx^/Mfn2 ^flx/flx^* mice, we conducted targeted metabolomics analysis on neural retinas collected from one-month-old mice. Three-dimentional principal component analysis (3D-PCA) of the metabolomics profiles revealed clear separation between *Rho-cre/Mfn1^flx/flx^/Mfn2 ^flx/flx^* groups and WT groups (Supplementary Fig. 3A). In total, 147 metabolites were identified, among which 23 metabolites were significantly upregulated and one was downregulated (p<0.05) (Fig. 4A and B). Our metabolomics analysis revealed significant alterations in metabolites associated with purine, pyrimidine, and lactate synthesis pathways, suggesting an overall upregulation of nucleotide biosynthesis in rod photoreceptor cells (Fig. 4B-D). Moreover, several amino acids such as glutamate and aspartate were significantly upregulated (Fig. 4B and Supplementary Fig. 3B).

**Figure 4.**
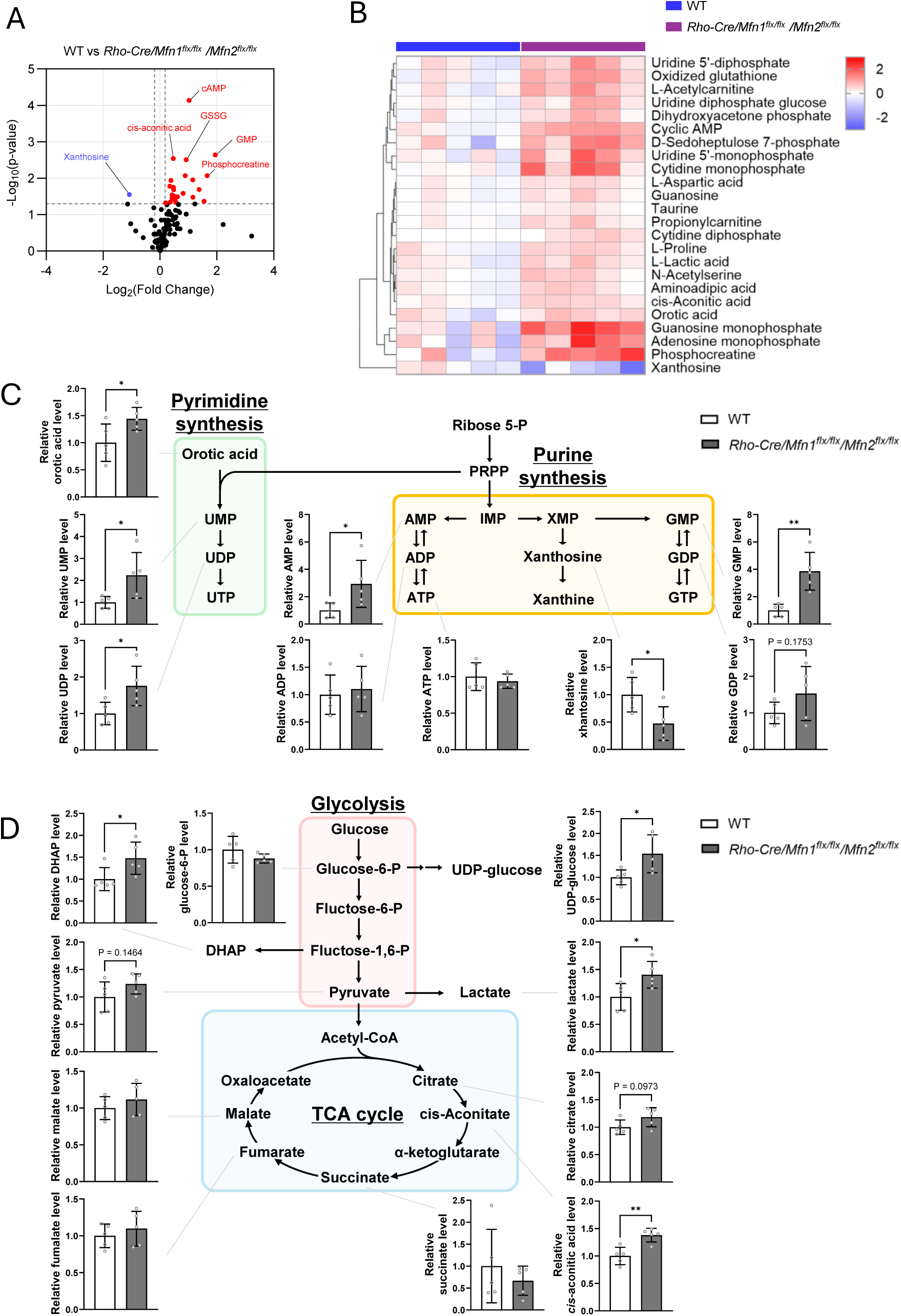
**Metabolic changes in the neural retina resulting from rod-specific ablation of *Mfn1* and *Mfn2*** (A) Volcano plot showing differentially changed metabolites in the neural retina of *Rho-Cre/Mfn1^flx/flx^/Mfn2^flx/flx^* mice versus WT mice. (B) Heatmap showing significantly changed metabolites in the neural retina of *Rho-Cre/Mfn1^flx/flx^/Mfn2^flx/flx^* mice versus WT mice (P<0.05). (C) Schematic diagram of pyrimidine and purine synthesis, and relative metabolite levels associated with nucleotide synthesis in *Rho-Cre/Mfn1^flx/flx^/Mfn2^flx/flx^* neural retina compared to WT neural retina. (D) Schematic diagram of glycolysis and TCA cycle, and relative metabolite levels associated with these pathways in *Rho-Cre/Mfn1^flx/flx^/Mfn2^flx/flx^* neural retina compared to WT neural retina. Data are presented as mean ± SD. Asterisks (*) indicate P < 0.05 significance by t-test. Five mice were used for each group in the study. Dots represent individual data points.

### Identified Changes in Protein Levels Associated with Pathways Presumed to Respond to Mitochondrial Fusion Defects

Our RNA-seq results indicated significant downregulation of genes involved in glycolysis (Fig. 3C). Rod photoreceptor cells require substantial energy for their function (21, 22) and mainly utilize glucose for aerobic glycolysis converting glucose to pyruvate, which is then converted to lactate by lactate dehydrogenase A (LDHA) (Fig. 5A) (5). Western blot analysis of proteins involved in glycolysis revealed a significant decrease in glyceraldehyde-3-phosphate dehydrogenase (GAPDH) expression, while pyruvate kinase M2 (PKM2) and LDHA levels remained unchanged (Fig. 5B). Subsequent to glycolysis, pyruvate is also converted to acetyl-CoA to replenish the TCA cycle followed by OXPHOS for energy production in photoreceptor cells (22), and therefore, we examined the expression of OXPHOS complex subunits. Notably, succinate dehydrogenase complex iron sulfur subunit B (SDHB), a component of Complex II that oxidizes FADH_2_ to FAD, was significantly decreased in *Rho-cre/Mfn1^flx/flx^/Mfn2 ^flx/flx^* mice (Fig. 5C). SDHA, another subunit of Complex II responsible for supplying SDHB with FADH_2_ synthesized via the TCA cycle, remained unchanged (Fig. 5D). In addition to the TCA cycle, mitochondrial β-oxidation serves as another source of FADH_2_ (23). The mitochondrial β-oxidation pathway begins with the uptake of acyl-CoA into mitochondria via carnitine-acylcarnitine translocase (CACT), followed by conversion to acyl-CoA by carnitine palmitoyl transferase 2 (CPT2) (Fig. 5E) (23). Western blot analysis revealed a significant reduction in CACT expression, while CPT2 levels remained unchanged (Fig. 5F), suggesting potentially lower flux through mitochondrial β-oxidation. In contrast, the expression of 70-kDa peroxisomal membrane protein (PMP70), a transporter involved in peroxisomal β-oxidation, and fatty acid synthase (FASN), an enzyme responsible for synthesizing fatty acids that serve as substrates for peroxisomal β-oxidation remained unchanged in *Rho-cre/Mfn1^flx/flx^/Mfn2 ^flx/flx^* mice (Supplementary Fig. 4), indicating that peroxisomal β-oxidation is not affected by mitochondrial changes.

**Figure 5.**
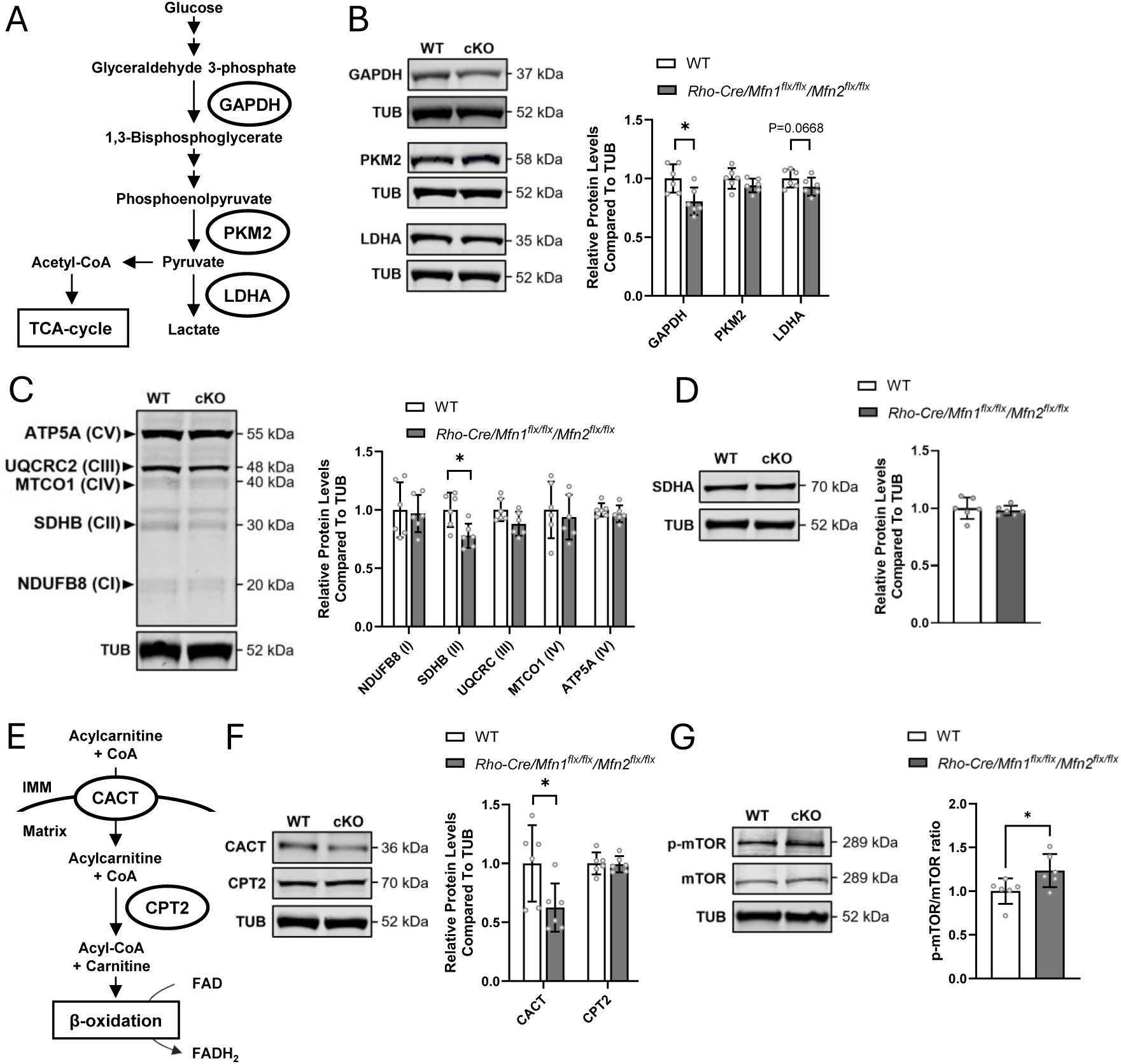
Identified changes in protein levels associated with pathways presumed to respond to mitochondrial fusion defects. (A) Schematic diagram of the glycolysis pathway of lactate synthesis from glucose through pyruvate in the cytosol of cells. (B) Western blot analysis of glyceraldehyde-3-phosphate dehydrogenase (GAPDH), pyruvate kinase M2 (PKM2), and lactate dehydrogenase (LDH), which are related to glycolysis pathway in the neural retina of *Rho-Cre/Mfn1^flx/flx^/Mfn2^flx/flx^* mice versus WT mice. (C) Western blot analysis of each subunit comprising the complexes (Complex I (CI): NADH dehydrogenase [ubiquinone] 1 beta subcomplex subunit 8 (NDUFB8), Complex II (CII): succinate dehydrogenase B (SDHB), Complex III (CIII): ubiquinol-cytochrome c reductase core protein 2 (UQCRC2), Complex IV (CIV): mitochondrially encoded cytochrome c oxidase I (MTCO1), Complex V (CV): ATP synthase F1 subunit alpha (ATP5A)) responsible for oxidative phosphorylation (OXPHOS). (D) Western blot analysis of another CII subunit, succinate dehydrogenase A (SDHA). (E) Schematic diagram of the substrate uptake and pathway toward mitochondrial β-oxidation. (F) Western blot analysis of carnitine-acylcarnitine translocase (CACT) and carnitine palmitoyl transferase II (CPT2), involved in mitochondrial β-oxidation. (G) Western blot analysis of mammalian target of rapamycin (mTOR) and phosphorylated-mTOR-S2448 (p-mTOR). Protein levels of p-mTOR were normalized by that of mTOR. Alpha-tubulin (TUB) served as the loading control for this Western blot experiments except for the result of p-mTOR. Data are presented as mean ± SD. Asterisks (*) indicates P < 0.05 significance following a significant difference detected by t-test. Six one-month-old mice were used in both groups in study. Dots represent individual data points. The protein size next to the immunoblot images denotes the size of the immunobands measured for this analysis.

mTORC1 serves as a central regulator of cellular homeostasis by integrating diverse cellular signals (24, 25). It is inhibited under low ATP conditions but becomes activated in response to amino acid and oxidative stress, both of which can be induced by ATP depletion (24, 25). Our RNA-seq analysis revealed that many of genes related to mTORC1 pathway are upregulated, along with amino acid metabolism-related genes in *Rho-cre/Mfn1^flx/flx^/Mfn2 ^flx/flx^* mice (Fig. 3C). Western blot analysis showed elevated levels of phosphorylated-mTOR (p-mTOR) levels (Fig. 5G), suggesting that mTOR activation may be triggered in *Rho-cre/Mfn1^flx/flx^/Mfn2 ^flx/flx^* mice indirectly in response to energy disruption.

Overall, these findings suggest that selective impairment of mitochondrial energy production through OXPHOS and β-oxidation occurs in *Rho-cre/Mfn1^flx/flx^/Mfn2 ^flx/flx^* mice, likely reducing the demand for pyruvate and driving metabolic reprogramming. mTOR activation likely occurs indirectly in response to abnormal energy production as a compensatory pathway to maintain homeostasis (Fig. 6).

**Figure 6.**
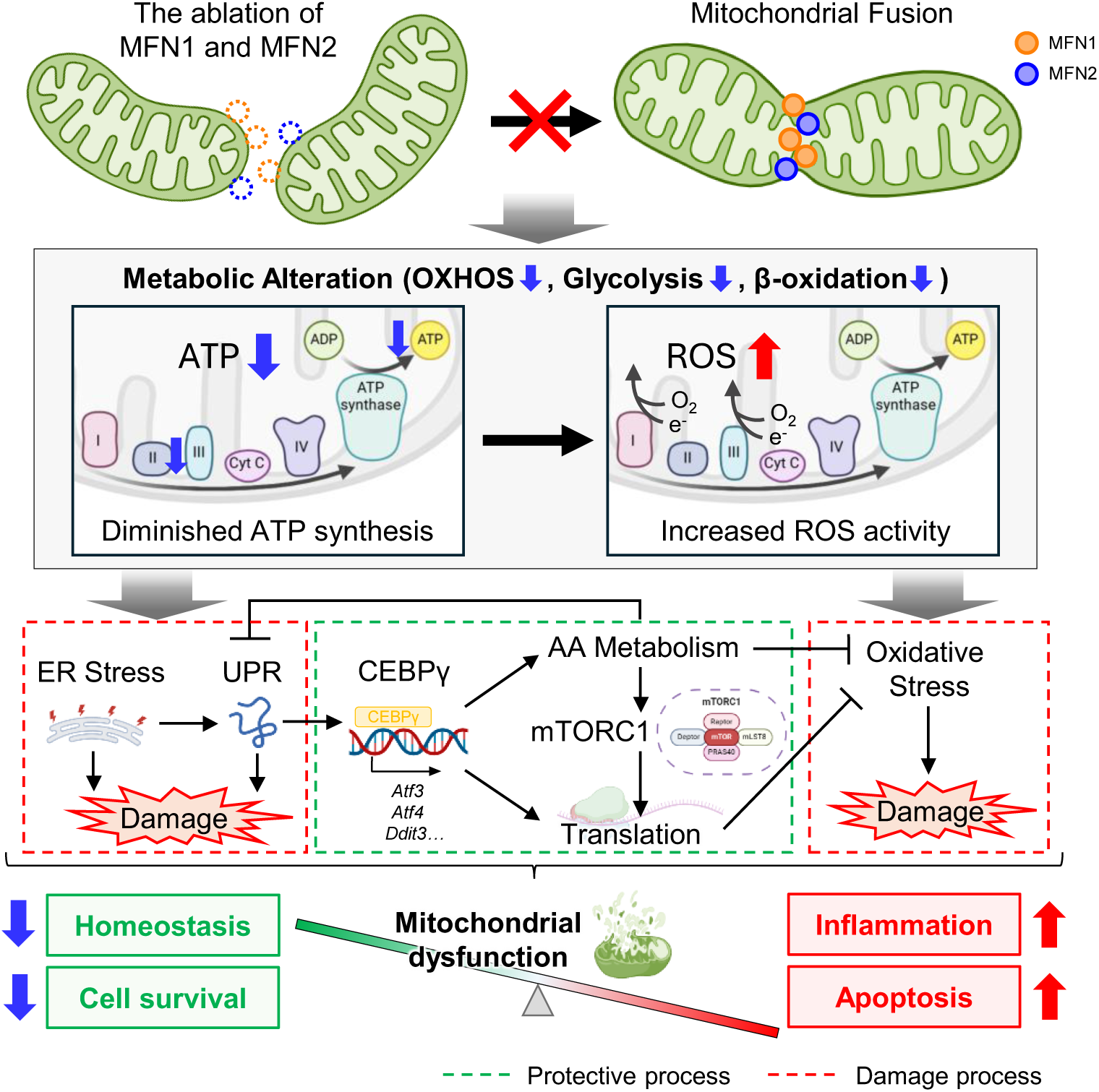
Graphical abstract summarizing the data presented in this study. Reduced energy production due to defective mitochondrial fusion in rod photoreceptor cells causes endoplasmic reticulum stress, UPR, and oxidative stress, in response to which the biosynthesis process via CEBPγ and mTOR pathways is activated. Despite the activation of these defense responses, mitochondrial fusion defects ultimately lead to photoreceptor cell death.

## Discussion

In this study, we demonstrated that MFN1 and MFN2 together are critical for establishing rod photoreceptor cell-specific mitochondrial morphology. Our findings further suggest that these specialized mitochondrial structures play a crucial role in sustaining proper energy production and overall maintenance of photoreceptor cells. Mitochondrial fusion defects were found to cause metabolic alterations due to reduced energy production, which in turn activate multiple cytotoxic pathways such as ER stress, UPR, and oxidative stress. In response, rod photoreceptor cells were also found to activate protective biosynthetic pathways such as AA synthesis and translation mediated by C/EBPγ and mTORC1 (Fig. 6) in an effort to maintain intracellular homeostasis.

### Mitofusins Regulate Mitochondrial Structures in Rod Photoreceptor Cells

Rod photoreceptor cells exhibit highly specialized mitochondrial architecture, characterized by a large, singular mitochondrion at the synaptic terminal and elongated mitochondria in the inner segment. Our findings suggest that mitochondrial fusion is essential for establishing these unique structures (Fig. 1A and C). Smaller mitochondria were observed in inner segments and synaptic terminals of *Rho-cre/Mfn1^flx/flx^/Mfn2 ^flx/flx^* mice, indicating that smaller mitochondria are trafficked to the synaptic terminal, where mitofusins mediate their subsequent fusion to form the distinctive morphologies. Interestingly, MFN2 has been implicated in mitochondrial trafficking into synaptic terminals in other neuronal cells (26, 27). However, our results indicate that mitofusins may not play a direct role in mitochondrial trafficking in rod photoreceptor cells, as smaller mitochondria were observed at their destinations without MFN2 or MFN1. This suggests a mitofusin-independent mechanism for mitochondrial trafficking in rod photoreceptor cells.

### Coordination Between MFN1 and MFN2

MFN1 and MFN2 are present on the outer mitochondrial membrane (OMM) and work together to regulate mitochondrial fusion (28–30). Our EM analysis revealed that depletion of both MFN1 and MFN2 caused mitochondrial fragmentation (Fig. 1A and C), suggesting that unique mitochondrial morphologies are developed through mitochondrial fusion mediated by both mitofusins. *Rho-cre/Mfn1^flx/+^/Mfn2 ^flx/flx^* mice and *Rho-cre/Mfn1^flx/flx^/Mfn2 ^+/+^* mice exhibited proper development of mitochondrial morphologies (Fig. 1A and C), indicating that MFN1 and MFN2 compensate for each other in terms of the formation of rod photoreceptor cell-specific mitochondrial morphologies. Consistent with our findings, a previous report showed that photoreceptor cell degeneration in MFN2-mutant mice (*MFN2^R94Q^*) is rescued by augmentation of MFN1 (*MFN2^R94Q^*:*MFN1*) (31). Mammalian MFN1 and MFN2 are very similar proteins with high homology (∼80%) (29, 30). Mitochondrial membrane fusion requires interaction of mitofusins with the C-terminal heptad repeat domain (HR2), and dimerization of the GTPase domain (29). Given the high degree of homology between MFN1 and MFN2 in both their GTPase and HR2 domains, it is plausible that they may functionally compensate for each other. However, *Rho-cre/Mfn1^flx/flx^/Mfn2 ^flx/+^* mice showed moderate mitochondrial fragmentation, while *Rho-cre/Mfn1^flx/+^/Mfn2 ^flx/flx^* mice displayed intact mitochondria (Fig. 1 A, B, D-F). This observation suggests that the functions of MFN1 and MFN2 are not completely redundant in mitochondrial fusion.

Several potential explanations may account for this observation. MFN1 has been shown to regulate inner mitochondrial membrane (IMM) fusion by controlling the expression levels of other mitochondrial dynamics proteins such as OPA1 and Fis1 (31). Therefore, in addition to affecting mitochondrial fusion through functions that are shared by MFN2 (and thus, can be compensated for by MFN2), MFN1 may also regulate mitochondrial fusion through distinct, MFN1-specific functions/mechanisms. Furthermore, it has been reported that MFN1 possesses eightfold higher GTPase activity compared to MFN2, suggesting a potential difference in their abilities to drive mitochondrial fusion (32). In addition, the protein levels of MFN1 and MFN2 may influence mitochondrial fusion, which may be different in rod photoreceptor cells. Finally, both MFN1 and MFN2 need to be recruited to the mitochondria to exert their functions in mitochondrial fusion. This recruitment to the OMM is mediated through interactions between their N-terminal mitochondrial targeting sequences and mitochondrial translocation complexes (28, 33). It is possible that differences exist between MFN1 and MFN2 in the process of mitochondrial recruitment.

### Defective Energy Production and Metabolic Adaptations

Mitochondrial dynamics play a critical role for optimal OXPHOS activity by allowing efficient transport and distribution of mitochondrial contents (34). Mitochondria contain OXPHOS complex subunits encoded by their small cyclic genome, and mitochondria fuse to maintain their functions (9). In *Rho-cre/Mfn1^flx/flx^/Mfn2 ^flx/flx^* mice, a reduction in SDHB, the complex II (CII) subunit involved in OXPHOS, was observed (Fig. 5C). CII plays a crucial role in oxidizing FADH_2_ for ATP production, thus its decrease in CII contributes to ATP deficiency (35). Recent studies have shown that decreased CII activity occurs in the brains of patients with neurodegenerative diseases such as Alzheimer’s, Parkinson’s, and Huntington’s disease (36–38). FADH_2_ is mainly produced via mitochondrial β-oxidation and glycolysis, which were both disturbed in *Rho-cre/Mfn1^flx/flx^/Mfn2 ^flx/flx^* mice (Fig. 3C, Fig. 5B and F). We revealed that CACT, responsible for acylcarnitine uptake into mitochondria as a first step of mitochondrial β-oxidation, was downregulated in *Rho-cre/Mfn1^flx/flx^/Mfn2 ^flx/flx^* mice (Fig. 5F), leading to the accumulation of acylcarnitines such as butyryl carnitine and propionyl carnitine as shown by our metabolomics analysis (Fig. 4B). Our results suggest that impaired mitochondrial fusion results in defective mitochondrial OXPHOS, and therefore, the demand for FADH_2_ produced by TCA cycle is reduced. Accordingly, the need for conversion of pyruvate to acetyl-CoA and its entry into the TCA cycle would be also reduced in these cells. In this scenario, pyruvate is more likely to be converted to lactate at the end of glycolysis, as a compensatory pathway. Increased lactate levels may lead to downregulation of GAPDH and hence glycolysis as the negative feedback to lower lactate levels (39–41). This could result in a decrease in pyruvate production and further decrease mitochondrial ATP production in *Rho-cre/Mfn1^flx/flx^/Mfn2 ^flx/flx^* mice. Besides our mouse model, it has been reported that rod photoreceptor cells lose normal function due to PKM2 deficiency (42). HK2, a glycolytic enzyme, is also known to contribute to rod photoreceptor survival from aging and stress responses (43, 44). While energy production through glycolysis, OXPHOS, and mitochondrial β-oxidation decreased, metabolomic analysis showed increased levels of L-proline and phosphocreatine (Fig. 4B), both of which are reported to maintain ATP synthesis in other cells (45–47). These results suggest that branching pathways for the ATP production are activated in *Rho-cre/Mfn1^flx/flx^/Mfn2 ^flx/flx^* mice.

### Induced Cellular Stress by Mitochondrial Abnormality and Compensatory Biological Reaction

Gene set enrichment analysis revealed significant upregulation of pathways related to ER stress and UPR in *Rho-cre/Mfn1^flx/flx^/Mfn2 ^flx/flx^* mice (Fig. 3A and B). ER stress is caused by various cellular stress, including energy deprivation and exposure to oxidative stress (48–50). Our results showed that ER stress can be induced by mitochondrial fusion defects, which was also observed in another model (51). Since ER stress accumulates damage to cells and causes apoptosis (52), various cytoprotective pathways exist to mitigate it. Biosynthesis of specific amino acids and cognate tRNA synthetases have been reported as biological processes to relieve ER stress (53). Our gene expression analyses suggest that *Rho-cre/Mfn1^flx/flx^/Mfn2 ^flx/flx^* mice exhibit an upregulation of amino acid metabolism and translation pathways including tRNA synthesis in response to cellular stress. This coordinated increase in protein translation is reminiscent of patterns observed in neurodegenerative diseases such as Huntington’s disease (54). Under ER stress and oxidative stress, translational reprogramming is mediated by stress response factors, including activating transcription factor 3 (ATF3), ATF4, and C/EBP homologous protein (CHOP) to preserve cellular homeostasis (55–58). Our RNA-seq data showed significantly elevated expression of these transcription factors (Fig. 3C), suggesting that translational reprograming is activated as an adaptive mechanism to counteract the cellular stress.

mTORC1 pathway, which is activated in *Rho-cre/Mfn1^flx/flx^/Mfn2 ^flx/flx^* mice (Fig. 3C and 5G), is known to regulate various biosynthesis processes (59, 60). mTORC1 acts in a cytoprotective manner by promoting mitochondrial biogenesis, nucleotide synthesis, and translation against cellular stress (24). Our metabolomics and RNA-seq analyses revealed that nucleotide synthesis and translation were activated in *Rho-cre/Mfn1^flx/flx^/Mfn2 ^flx/flx^* mice (Fig. 3B, 4B and C). Activation of mTORC1 is induced by AA metabolism and oxidative stress, both of which are known biological processes that occur in mitochondrial stress (24, 25, 61), and consistent with our results. Additionally, metabolomics analysis identified upregulation of other metabolites increasing mTORC1 activity, including DHAP (Fig. 4B and D) (62, 63). Reduced glycolysis (Fig. 3C and 5B) may lead to increased conversion of GAP to DHAP, further driving mTORC1 activation.

OXPHOS dysfunction and mitochondrial stress can enhance reactive oxygen species (ROS) production, further exacerbating cellular stress. The elevated glutathione disulfide (GSSG) levels in *Rho-cre/Mfn1^flx/flx^/Mfn2 ^flx/flx^* mice provide supporting evidence for this. Oxidative stress in mitochondria has been reported to cause ER stress (64). Various defense mechanisms against oxidative stress exist in cells (65). Glutathione (GSH), made of amino acids, oxidizes itself to GSSG, which contributes to neutralization of oxidative stress (65, 66). In *Rho-cre/Mfn1^flx/flx^/Mfn2 ^flx/flx^* mice, the accumulation of GSSG in the neural retina indicates that a defense mechanism against oxidative stress is at work (Fig. 4B). In addition to GSH, we believe that the translational activation mentioned above also contributes to the maintenance of the redox status (25, 58, 67). While various biological defense responses are induced to resist intracellular stress, our study indicated that chronic mitochondrial fusion failure appeared to cause more damage than bioprotective effects, ultimately leading to rod photoreceptor cell death.

### Limitation of our study

In our study, we used protein abundance analysis to identify changes in the expression of mitochondrial metabolism related proteins in response to mitochondrial morphological dysfunction. We found that failure of mitochondrial fusion leads to decreased expression of several proteins involved in OXPHOS, glycolysis, and mitochondrial β-oxidation. However, the lysates used in these studies contain proteins from all cells in the neural retina. Therefore, our analysis may underestimate the changes in protein expression that occur within rod photoreceptor cells, and more genes and proteins may be altered in these cells. In future studies, analyses employing a single-cell approach specific to rod photoreceptor cells would be very beneficial as it would allow a deeper exploration of other key players that respond to defects in mitochondrial dynamics.

## Conclusion

In conclusion, using photoreceptor cells as a model, our study showed that cell-type specific mitochondrial structures are critical for cell-type optimized energy production. We showed that unique mitochondrial morphologies in rod photoreceptor cells are formed and regulated by mitochondrial fusion. Furthermore, mitochondrial fusion plays a crucial role in maintaining energy production and function in rod photoreceptor cells, and its impairment severely damage rod photoreceptor cells. Our present study identified cellular stress pathways and compensatory cytoprotective pathways in response to mitochondrial dynamics failures. Mitochondrial dysfunction has been increasingly recognized as a key contributor to aging and neurodegenerative diseases (68, 69). The pathways identified in our study provide valuable insights and potential entry points for further investigation into mitochondrial dysfunction-associated neuronal disorders. A deeper understanding of how mitochondrial fusion supports cellular homeostasis will be pivotal in elucidating its role in retinal diseases.

## Materials and Methods

### Animals

*Rho-Cre* mice (B6.Cg-*Pde6b^+^* Tg[Rho-icre]1Ck/Boc [JAX stock #015850]) (17), *Mfn1* floxed mice (*Mfn1^flx/flx^* ; B6.129[Cg]-*Mfn1^tm2Dcc^*/J [JAX stock #026401]) (17), and *Mfn2* floxed mice (*Mfn2 ^flx/flx^* ; B6.129[Cg]-*Mfn2^tm3Dcc^*/J [JAX stock #026525]) (17) were purchased from The Jackson Laboratory. All the strains were congenic on the C57BL/6J background and tested negative for *Pde6b^rd1^* and *Crb1^rd8^* mutations. They were bred together to generate Rho-icre*/Mfn1^flx/flx^/Mfn2 ^flx/flx^* mice on the C57BL/6J background used in this study. WT mice on the C57BL/6 J background were used as controls for these experiments. All animals were housed in the same animal facility at the University of Wisconsin-Madison under the same environmental conditions. Both male and female mice that were one month and three months of age were used in this study. All experiments performed in this study were in accordance with the National Institute of Health Guide for the Care and Use of Laboratory Animals and authorized by the Animal Care and Use Committee at the University of Wisconsin-Madison. The results and methods in this study are reported in accordance with the ARRIVE guidelines.

### Electron Microscopy

Eyes were fixed with 2% paraformaldehyde (PFA) and 2% glutaraldehyde and submitted to the Electron Microscope Core at the University of Wisconsin-Madison for transmission EM processing as previously described (70–72). Eye sections were mounted on a 400-mesh thin bar grid, and images were collected where the grid bars intersected the neural retinas using a Phillips CM120 STEM microscope (FEI Company, Hillsboro, OR, USA) at 8,800X magnification. Mitochondria numbers were counted using NIH’s ImageJ software.

### Immunohistochemistry

Eyes were punctured with a needle in the cornea and fixed with 4% paraformaldehyde (PFA) for 30 minutes at room temperature. Then the cornea and lens were removed, and neural retinas were separated from the eyecups. Neural retinas were blocked in 10% normal donkey serum for 30 minutes at room temperature. Next, they were incubated overnight with the 1:50 diluted primary antibody against TOMM20 (#sc-17764, Santa Cruz Biotechnology, Hercules, TX, USA) at 4°C with slow shaking. They were rinsed in PBS, and incubated with a 1:250 diluted Donkey Anti-Mouse IgG H&L (Alexa Fluor® 488) (#ab150105, Abcam, Cambridge, UK) for 45 min at room temperature. Before mounting, four small incisions were made to permit flattening of the retina. Retinal whole mounts were imaged to visualize mitochondria in the inner segment using SoRa/W1 Spinning Disk Microscope (Nikon, Tokyo, Japan) at a 100X magnification.

### Histological analysis

Eyes were fixed with 2% PFA and 2% glutaraldehyde overnight. Eyes were then rinsed with PBS and embedded in paraffin. Samples were submitted to Translational Research Initiatives in Pathology (TRIP) core at the University of Wisconsin-Madison for processing and sectioning. Six μm sections were cut on a RM 2135 microtome (Leica Microsystems, Wetzlar, Germany). Paraffi sections were stained with hematoxylin and eosin (H&E) using stand protocols to visualize retinal layers and imaged using an Axio Imager 2 microscope (Carl Zeiss MicroImaging, NY, USA) at a 40X magnification.

### Bulk RNA-sequencing

Neural retinas were collected and pooled from individual one-month-old WT and *Rho-cre/Mfn1^flx/flx^/Mfn2 ^flx/flx^* mice between 11:00 AM and 1:00 PM. Samples were flash frozen and then submitted to GENEWIZ from Azenta Life Scienses (South Plainfield, NJ, USA) for processing. Total RNA was extracted from neural retinas with an RNeasy Plus Universal Mini kit (Qiagen, Hilden, Germany) following the Manufacturer’s protocols. Total RNA samples were quantified using a Qubit 2.0 Fluorometer (Life Technologies, Carlsbad, CA, USA), and RNA integrity was examined using a TapeStation 4200 (Agilent Technologies, Palo Alto, CA, USA). RNA sequencing libraries were prepared using the NEBNext Ultra RNA Library Prep Kit for Illumina (NEB, Ipswich, MA, USA) following manufacturer’s instructions. Briefly, messenger RNAs were first enriched with Oligo(dT) beads. Enriched mRNAs were fragmented for 15 min at 94 °C. First strand and second strand cDNAs were subsequently synthesized. cDNA fragments were end repaired and adenylated at 3’ends, and universal adapters were ligated to cDNA fragments, followed by index addition and library enrichment by limited-cycle PCR. The sequencing libraries were validated on the Agilent TapeStation and quantified by using Qubit 2.0 Fluorometer as well as by quantitative PCR (KAPA Biosystems, Wilmington, MA, USA). The sequencing libraries were clustered on a flowcell. After clustering, the flowcell was loaded on the Illumina HiSeq instrument (4000 or equivalent) according to manufacturer’s instructions. The samples were sequenced using a 2 × 150 bp Paired End (PE) configuration. Image analysis and base calling were conducted by the HiSeq Control Software (HCS). Raw sequence data (.bcl files) generated from Illumina HiSeq were converted into fastq files and de-multiplexed using Illumina’s bcl2fastq 2.17 software. The RNA-Seq raw sequence files from this study are available on the Gene Expression Omnibus (GEO), accession number GSE297370.

### RNA-sequencing analysis

Gene expression read counts were analysed using NetworkAnalyst 3.0 (73). *M. musculus* organism was selected with bulk sequencing analysis workflow. Quality control step involved filtering genes with very high variance across samples. Genes were ranked based on variance and those genes which ranked in the bottom 15% of the percentile were filtered out. Low abundance genes below a threshold were also filtered out. Data was normalized using log 2 counts per million normalization method. Differential gene expression analysis was performed using EdgeR (74). Gene set enrichment analysis was performed using WebGestalt (75)and different functional databases including Gene Ontology, KEGG, and WikiPathways were used for analysis. To further restrict the number of gene sets due to overlap of the genes, affinity cluster algorithm (76) was applied. Signalling pathway analysis was conducted on differentially expressed genes using the SIGNOR 2.0 database (77).

### Metabolomics

Neural retinal tissues were isolated from mice and stored at −80 °C. These samples were submitted to Metabolomics Core Resource Laboratory at New York University. Each tissue sample was then weighed and transferred into a bead blaster tube on dry ice. Prior to extraction, 80% methanol in water containing the internal standard (AA standard) was placed on dry ice for approximately 15 minutes.

For homogenization and extraction, 100 μL of glass beads was added to each bead blaster tube containing the tissue sample, followed by the addition of 80% methanol in water with the AA standard to achieve a final tissue concentration of 10 mg/mL. The samples were homogenized using a bead blaster for 10 cycles, with each cycle consisting of 30 seconds on and 30 seconds off. Following homogenization, the samples were centrifuged at 21,000 xg for 3 minutes. A total of 450 μL of the supernatant was then collected from each. These collected supernatants were dried down using a SpeedVac, after which the dried sample was reconstituted in 50 μL of mass spectrometry-grade (MA grade) water. The reconstituted sample was sonicated for 2 minutes and subsequently centrifuged at 21,000 xg for 3 minutes. Finally, 20 μL of the processed sample was transferred into a glass insert within a 2 mL glass vial for analysis. Samples were analyzed with the hybrid LCMS assay after scaling the metabolite extraction to a measured aliquot (10mg/mL) for each sample and metabolites were quantified. Overall, coverage of the library was 147 metabolites being detected. The resulting data were analyzed by principal components analysis (PCA), visualizing clusters, volcano plots, and other statistical comparisons. Data files have been uploaded to MetaboLights database (ID: MTBLS12512), http://www.ebi.ac.uk/metabolights/.

### Western blot analysis

Tissues were isolated from mice and stored at −80 °C. Neural retina lysates were homogenized using a Bel-Art Homogenizer system motor in RIPA buffer (#P189901, Thermo Fisher Scientific, Waltham, MA) containing protease inhibitors (#11836170001, Thermo Fisher Scientific, Waltham, MA), respectively. Protein concentrations were quantified using a BCA Protein Assay Kit (#P123228, Thermo Fisher Scientific, Waltham, MA). Equal protein amounts were aliquoted, reduced using XT Reducing Agent (#1610792, Biorad, Hercules, CA) for seven minutes at 105 °C, and loaded onto 10% Bis-Tris Criterion XT gels (#3450112, Biorad, Hercules, CA) in MOPS buffer (#1610788, Biorad, Hercules, CA) and transferred to nitrocellulose membranes (#102673-324, Biorad, Hercules, CA) or Immun-Blot PVDF membranes for Protein Blotting (#1620177, Biorad, Hercules, CA). Membranes were blocked with milk or EveryBlot Blocking Buffer (#12010020, Biorad, Hercules, CA), and probed overnight with their respective primary antibody at 4 °C. The primary antibodies and their dilutions used in this study can be found in Supplementary Table 1. Blots were washed with TBST buffer the next day and incubated with their corresponding secondary antibody. Secondary antibodies used in this study included donkey anti-rabbit IgG 680RD (#926-68073, LI-COR), donkey anti-rabbit IgG 800CW (#926-32213, LI-COR), donkey anti-goat IgG 680RD (#926-68074, LI-COR), goat anti-mouse IgG1 800CW (#926-32350, LI-COR), goat anti-mouse IgG2a 800CW (#926-32351, LI-COR), and goat anti-mouse IgM 800CW (#925-32280, LI-COR). Blots were washed again with TBST and imaged using the Odyssey Imaging System (LI-COR Biosciences, Lincoln, NE) and analyzed using NIH’s ImageJ (Bestheda, MD). Blots were stripped with Newblot Stripping Buffer (LI-COR Biosciences, Lincoln, NE) according to the manufacturer’s protocol and re-probed with another primary antibody in this study. All immunobands were normalized to the loading control on their respective immunoblot.

### Statistical Analysis

All statistical tests were performed using Prism Software (GraphPad, San Diego, CA). Significance of the difference between groups was calculated by unpaired Student’s two-tailed t test, for experiments comparing two groups, and one-way or two-way analyses of variance (ANOVA) with the Bonferroni-Dunn multiple comparison posttest for experiments comparing three or more groups using *p<0.05, **p<0.01, ***p<0.001. ****p<0.0001. All data are presented as the mean ± the standard deviation(s.d.) of three or more independent experiments, with three or more replicates per condition per experiment. P < 0.05 was considered to be statistically significant.

## Supporting information

Supprementary Figure

## Author Contributions

Conceptualization – ML, RH, VB, TT, AI; Data curation – ML, RH, PG, VB, TT; Formal analysis – ML, RH, PG, TT, VB; Funding Acquisition – ML, AI; Investigation – ML, RH, PG, VB, TT; Methodology – ML, RH, PG, VB, TT, SI, AI; Project Administration – ML, RH, PG, SI, AI; Resources – SI, AI; Supervision – SI, AI; Validation – ML, RH, PG, VB, TT, SI, AI; Visualization – ML, RH, PG, TT; Writing-Original Draft – RH, SI, AI; Writing-Review and Editing – ML, RH, PG, VB, SI, TT, AI

## Classification

Major classification: Retina, Genetics, Minor classification: Photoreceptors, Mitochondrial dynamics

## Acknowledgement and funding sources

The authors would like to thank Toshi Kinoshita and the University of Wisconsin (UW) Translational Research Initiatives in Pathology laboratory (TRIP), supported by the UW Department of Pathology and Laboratory Medicine, UWCCC (P30 CA014520), and the Office of The Director-NIH (S10OD023526) for the use of facilities and services, and Randall Massey and the University of Wisconsin Electron Microscope Core for tissue processing, sectioning, and assistance for this study. The authors would like to thank Department of Biochemistry at the University of Wisconsin-Madison for imaging using SoRa/W1 Spinning Disk Microscope. The authors want to thank GENEWIZ from Azenta Life Scienses for their assistance with our RNA-seq analysis. The authors would also like to extend their gratitude to Dr. Drew Jones, and the New York University Langone Medical Center Metabolomics Core Resource Laboratory for their time and efforts in optimizing the protocols for our metabolomics experiments.

This work was supported by grants from the National Eye Institute (R01EY022086 and R01EY036383 to A. Ikeda; P30EY016665 to the Department of Ophthalmology and Visual Sciences at the University of Wisconsin-Madison; F32EY032766 to M. Landowski), and Timothy William Trout Chairmanship (A. Ikeda).

## Abbreviations

3D-PCA: 3-dimentional principal component analysis
AA: amino acids
ADP: adenosine diphosphate
AMP: adenosine monophosphate
ATF3: activating transcription factor 3
ATF4: activating transcription factor 4
ATP: adenosine triphosphate
ATP5A: ATP synthase F1 subunit alpha
AvgExpr: average expression
CI: Complex I
CII: Complex II
CIII: Complex III
CIV: Complex IV
CV: Complex V
C/EBPγ: CCAAT/enhancer-binding protein Gamma
CACT: carnitine-acylcarnitine translocase
Cho: choroid
cKO: conditional knockout
CoA: coenzyme A
Cone: cone photoreceptors
CHOP: C/EBP homologous protein
CPT2: carnitine palmitoyl transferase II
DHAP: dihydroxyacetone phosphate
EM: electron microscopy
ER: endoplasmic reticulum
FAD: Flavin adenine dinucleotide
FASN: fatty acid synthase
FC: fold change
FDR: false discovery rate
Fis1 Fission: Mitochondrial 1
GAP: glyceraldehyde-3-phosphate
GAPDH: glyceraldehyde-3-phosphate dehydrogenase
GDP: guanosine diphosphate
GMP: guanosine monophosphate
GSEA: gene set enrichment analysis
GSH: glutathione
GSSG: glutathione disulfide
GTP: guanosine triphosphate
H&E: hematoxylin and eosin
HR2: C-terminal heptad repeat domain
IMM: inner mitochondrial membrane
IMP: inosine monophosphate
INL: inner nuclear layer
IPL: inner plexiform layer
IS: inner segments
LDH: lactate dehydrogenase
Mac: macrophages
Mfn1: Mitofusin 1
Mfn2: Mitofusin 2
Mfns: Mitofusins
MTCO1: mitochondrially encoded cytochrome c oxidase I
mTOR: mammalian target of rapamycin
NDUFB8: NADH dehydrogenase [ubiquinone] 1 beta subcomplex subunit 8
NES: normalized enrichment score
OligoDend: oligodendrocytes
OMM: outer mitochondrial membrane
ONL: outer nuclear layer
ONLT: outer nuclear layer thickness
OPA: OPA1 mitochondrial dynamin like GTPase
OPL: outer plexiform layer
OS: outer segments
OXPHOS: oxidative phosphorylation
PFA: paraformaldehyde
PKM2: pyruvate kinase M2
PMP70: 70-kDa peroxisomal membrane protein
p-mTOR: phosphorylated mammalian target of rapamycin
PRPP: phosphoribosyl pyrophosphate
RGC: retinal ganglion cells
Rho: rhodopsin
RNA-seq: RNA-sequencing
Rod: rod photoreceptors
ROS: reactive oxygen species
RPE: retinal pigment epithelium
SDHA: succinate dehydrogenase A
SDHB: succinate dehydrogenase B
TCA: tricarboxylic acid
TOMM20: translocase of outer mitochondrial membrane 20
tRNA: transfer RNA
TUB: alpha-tubulin
UDP: uridine diphosphate
UMP: uridine monophosphate
UPR: unfold protein response
UTP: uridine triphosphate
UQCRC2: ubiquinol-cytochrome c reductase core protein 2
XMP: xanthosine monophosphate

## Data Availability Statement

Raw RNA-seq and metabolomics data have been uploaded to the GEO and MetaboLights database. The raw RNA sequencing data generated in this study have been deposited in the Gene Expression Omnibus (GEO) under accession number GSE297370. The metabolomics data have been deposited in the MetaboLights database under accession ID MTBLS12512 and are accessible at http://www.ebi.ac.uk/metabolights/.

