## Supplementary material for "Mitochondrial fusion controls the development of specialized mitochondrial structure and metabolism in rod photoreceptor cells": Supprementary Figure

**This PDF file includes:**

Figures S1 to S4
Tables S1

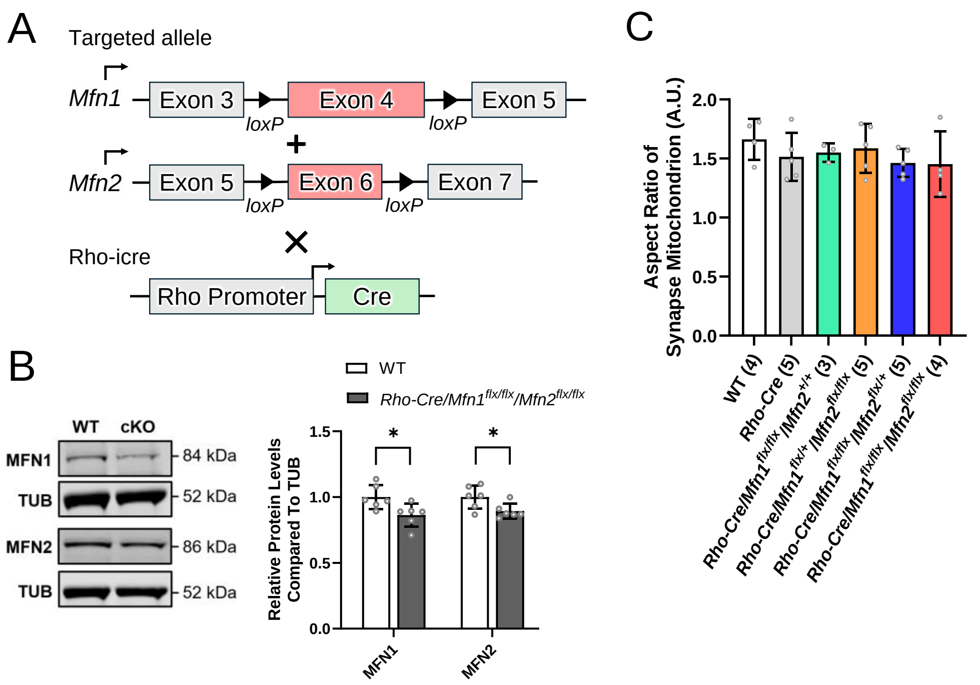


**Supplementary Figure 1. Combined ablation of *Mfn1* and *Mfn2* specifically in rod photoreceptor cells.**
(A) Schematic diagram of Cre-Lox strategy for rod photoreceptor cell-specific ablation of mitofusin1 (*Mfn1*) and mitofusin2 (*Mfn2*). (B) Western blot analysis of MFN1 and MFN2 using neural retinas from one-month-old WT and *Rho-Cre/Mfn1*$^{flx/flx}$/*Mfn2*$^{flx/flx}$ mice. Alpha-tubulin (TUB) served as the loading control. Data are presented as mean ± SD. Asterisks (*) indicates $P < 0.05$ significance by t-test. Six one-month-old mice were used for each group. Dots represent individual data points. The protein size next to the immunoblot images denotes the size of the immunoband measured for this analysis. (C) Quantification of the aspect ratio of mitochondria in the photoreceptor cell synaptic terminal. Number in the parenthesis denotes the number of mice used in the study. Data is presented as mean +/- SD and analyzed by two-way ANOVA with post-hoc Tukey's test.

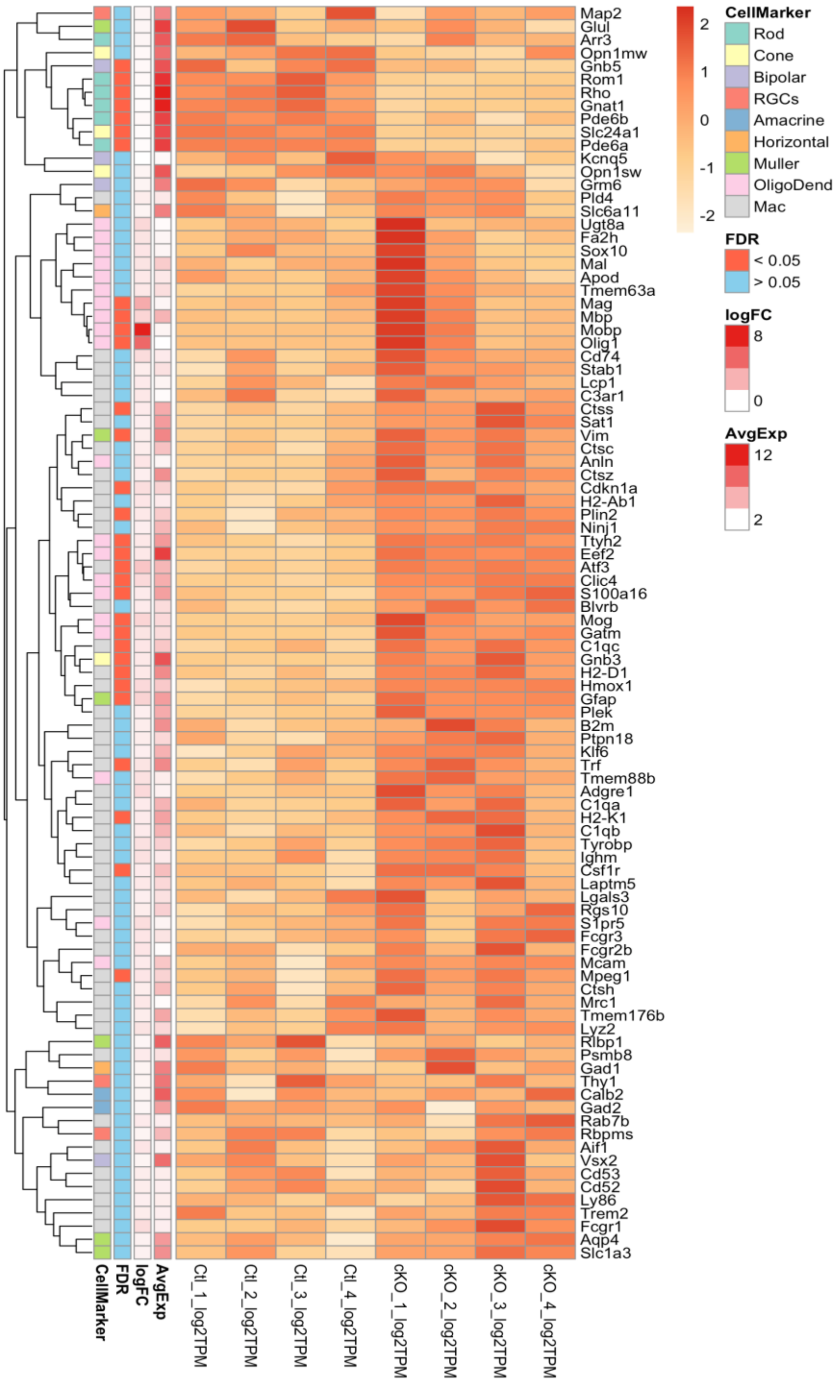


**Supplementary Figure 2. Gene expression analysis of specific markers for retinal cell types in mice with rod-specific ablation of *Mfn1* and *Mfn2***

Heatmap showing genes encoding specific markers for cells comprising the retina: rod photoreceptor cells (Rod), cone photoreceptor cells (Cone), bipolar cells, retinal ganglion cells (RGC), amacrine cells, horizontal cells, Müller cells, oligodendrocyte cells (OligoDend), macrophages (Mac). For each gene, false discovery rate (FDR), logFC, and average expression (AvgExpr) are shown in the left column.

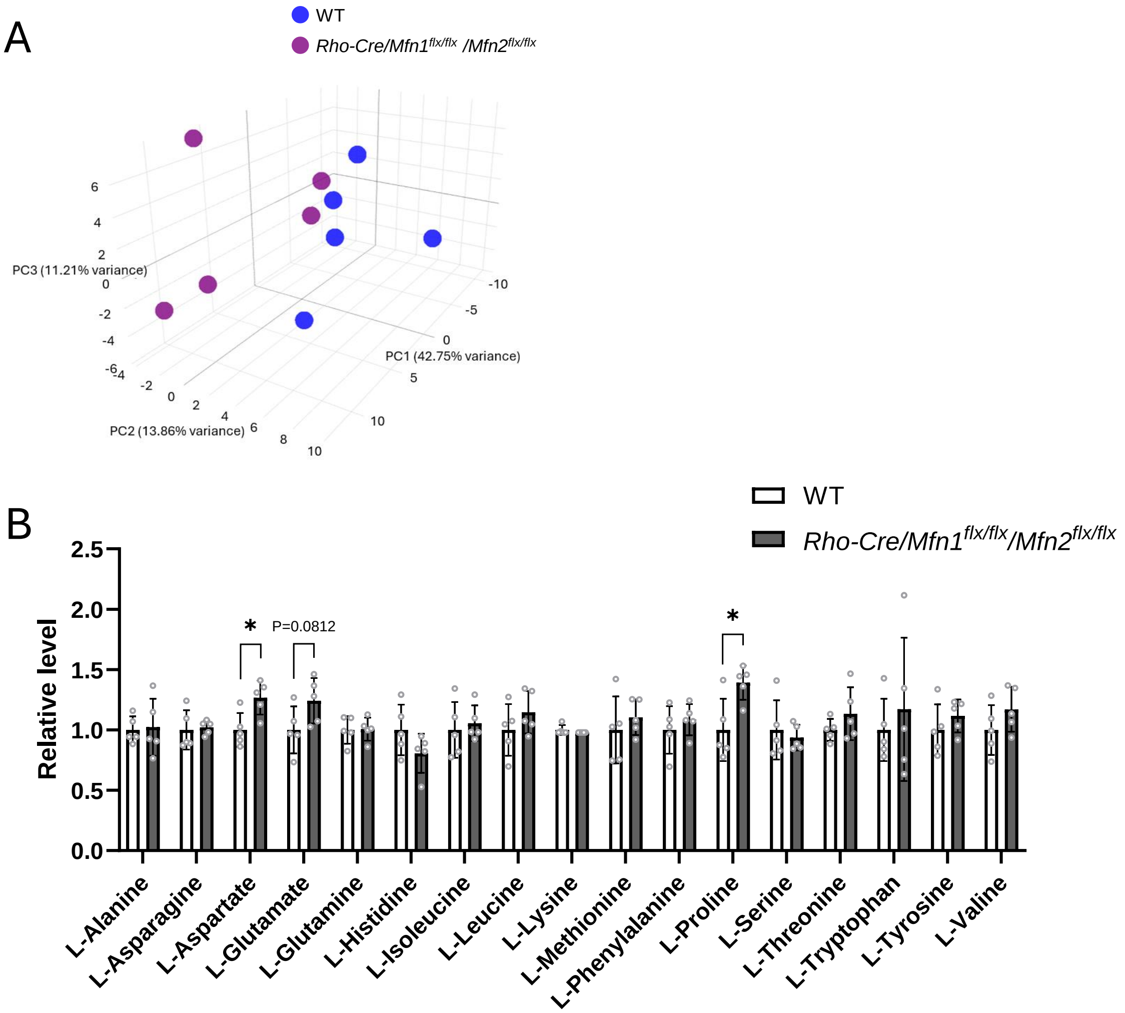


**Supplementary Figure 3. Three-dimensional principal component analysis and analysis of amino acid levels in the retina of mice with rod-specific ablation of *Mfn1* and *Mfn2***
(A) Three-dimensional principal component analysis of specialized metabolite components in *Rho-Cre/Mfn1*$^{flx/flx}$/*Mfn2*$^{flx/flx}$ neural retinas compared to WT neural retinas. (B) Relative amino acid levels in the neural retina of *Rho-Cre/Mfn1*$^{flx/flx}$/*Mfn2*$^{flx/flx}$ mice compared to WT mice. Data are presented as mean ± SD. Asterisks (*) indicate $P < 0.05$ significance by t-test. Five mice were used for each group in the study. Dots represent individual data points.

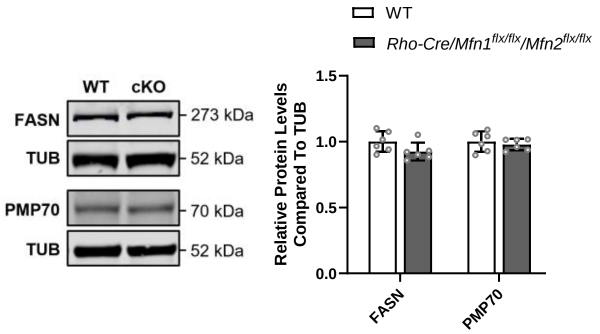


**Supplementary Figure 4. Levels of proteins related to peroxisomal β-oxidation in the retina of mice with rod-specific ablation of *Mfn1* and *Mfn2***

Western blot analysis of 70-kDa peroxisomal membrane protein (PMP70) and fatty acid synthase (FASN), related to peroxisomal β-oxidation in neural retinas from one-month-old WT and *Rho-Cre/Mfn1*$^{flx/flx}$*/Mfn2*$^{flx/flx}$ mice. Alpha-tubulin (TUB) served as the loading control. Data are presented as mean ± SD. Asterisks (*) indicates $P < 0.05$ significance following a significant difference detected by t-test. Six one-month-old mice were used for each group in the study. Dots represent individual data points. The protein size next to the immunoblot images denotes the size of the immunoband measured for this analysis.

**Table 1. Primary antibodies used in western blot analyses**

| Primary antibody (dilution) | Species | Catalog No. | Manufacturer |
|---|---|---|---|
| MFN1 (1:1000) | Rabbit | 10089-446 | Proteintech |
| MFN2 (1:1000) | Rabbit | 9482 | Cell Signaling Technology |
| mTOR (1:750) | Rabbit | 2983 | Cell Signaling Technology |
| p-mTOR (1:500) | Rabbit | 2971S | Cell Signaling Technology |
| GAPDH (1:2000) | Goat | PLA0302 | Sigma-Aldrich |
| PKM2 (1:1000) | Rabbit | 15822-1-AP | Proteintech |
| LDHA (1:1000) | Rabbit | 19987-1-AP | Proteintech |
| OXPHOS (1:1000) | Mouse | AB110413 | Abcam |
| SDHA (1:1000) | Rabbit | 11998 | Cell Signaling Technology |
| CACT (1:1000) | Rabbit | 19363-1-AP | Proteintech |
| CPT2 (1:1000) | Rabbit | 26555-1-AP | Proteintech |
| FASN (1:1000) | Rabbit | ab22759 | Abcam |
| PMP70 (1:1000) | Rabbit | ab3421 | Abcam |
| TUB (1:2000) | Mouse | 3873 | Cell Signaling Technology |
